# Isolation and *in vitro* culture of primary *pia mater* cells

**DOI:** 10.1101/2021.02.01.429222

**Authors:** Fadi Saadeh, Jan Remsik, Camille Derderian, Yudan Chi, Adrienne Boire

**Affiliations:** Human Oncology and Pathogenesis Program, 1275 York Avenue, New York, NY 10065; Department of Neurology, 1275 York Avenue, New York, NY 10065; Brain Tumor Center, 1275 York Avenue, New York, NY 10065; Memorial Sloan Kettering Cancer Center, 1275 York Avenue, New York, NY 10065

## Abstract

The meninges remain an unexplored area of neurobiology. These structures play host to dozens of morbid pathologies. This protocol provides a reliable way to identify and isolate pial cells from mice using robust markers of pial identity in mouse and human tissues. We describe a protocol for the extraction of *pia mater* cells from mice and their culture as primary cells *in vitro*. Using an array of transcriptomic, histological, and flow cytometric analyses, we identified Icam1 and Slc38a2 as two novel *pia mater* markers *in vitro* and *in vivo*. Our results confirm the fibroblastoid nature of pial cells and their ability to form a sheet-like layer that covers the brain parenchyma. To our knowledge, this is the first published protocol for the isolation, tissue culture, and marker identification of pial cells from mice. These findings will enable researchers in CNS barriers to describe pial cell functions in both health and disease.

## Introduction

The meninges are complex connective tissue structures that surround the brain and spinal cord. Meninges are anatomically divided into the *dura mater* and leptomeninges which consists of the *arachnoid mater* and *pia*. Embryologically, meningeal layers are derived from mesenchymal tissue that surrounds the neural tube during development. The *dura mater* is a well innervated, highly vascularized collagenous membrane that also contains lymphatics ^1,2^. Lying just below the subdural space, the *arachnoid mater*, forming a multilayer membrane covers the cerebrospinal fluid-filled subarachnoid space, creating a cellular barrier through its tight junctions ^3,4^. Proximal to the brain parenchyma, the *pia mater*, a one to two cell-layer membrane, covers the brain and spinal cord. Intimately associated with nervous tissue, the *pia* extends into sulci and fissures, delves deep into the parenchyma reflecting on subarachnoid vessels and to join the membranous ventricular lining.

The leptomeninges host a wide variety of physiologic and pathologic processes, including meningitis, immunoinflammatory disorders, leptomeningeal metastasis, and others. However, our current understanding of the pathophysiology of these processes is largely limited to events occurring in the CSF. The role of *pia* remains vastly unexplored. Many studies highlight the importance of the leptomeninges in influencing major brain centers and cortical layer formation during development ^5,6^, but it is unclear how these cells directly interact with the parenchyma. Interestingly, pathology affecting the parenchyma, being infectious or non-infectious, is almost always preceded by meningeal inflammation ^7–10^. A recent study identifies S100a6 as a marker for pial cells in embryonic meningeal tissue ^11^. However, no studies address the durability of this marker or presence of any other marker for *pia mater* cells in adult mice or humans. Interest in the leptomeninges has been on the rise due to its implication in metastases from several tumors, inflicting dismal prognoses. In order to study this essential component of this system, examination of normal *pia mater* cells is crucial. Described here is a detailed method for isolation and primary culture of *pia mater* cells from adult mice.

## Results

### Culture of pial cells

The *pia mater* isolated from mice contains a variety of cell types including immune, endothelial and other cells. The brain-specific marker of adult pial cells in not yet known. Exclusion of known cell types (CD45+, CD31+) allowed us to obtain a sample enriched in pial cells (**Figure 2a, b**). Pial cells acquired a fibroblast-like morphology *in vitro*, with flat oval nuclei and spindle-shaped cell body (**Figure 1b**). To assess the feasibility of maintaining pial cells in culture, we assessed the viability of pial cells with *in vitro* passaging and repetitive freeze-thaw cycles. Given the primary nature of these cells, viability of pial cells decreased by about 5% with every passage reaching 42% viability after 10 passages (**Supplementary Figure 1a**). Viability further decreased with increasing numbers of freeze-thaw cycles pial cells were subjected to (**Supplementary Figure 1b**). Pial cells also display increased cellular senescence with *in vitro* passaging shown, by accumulation of *Cdkn2a* expression with increased passage numbers (**Supplementary Figure 1c**).

**Figure 1:**
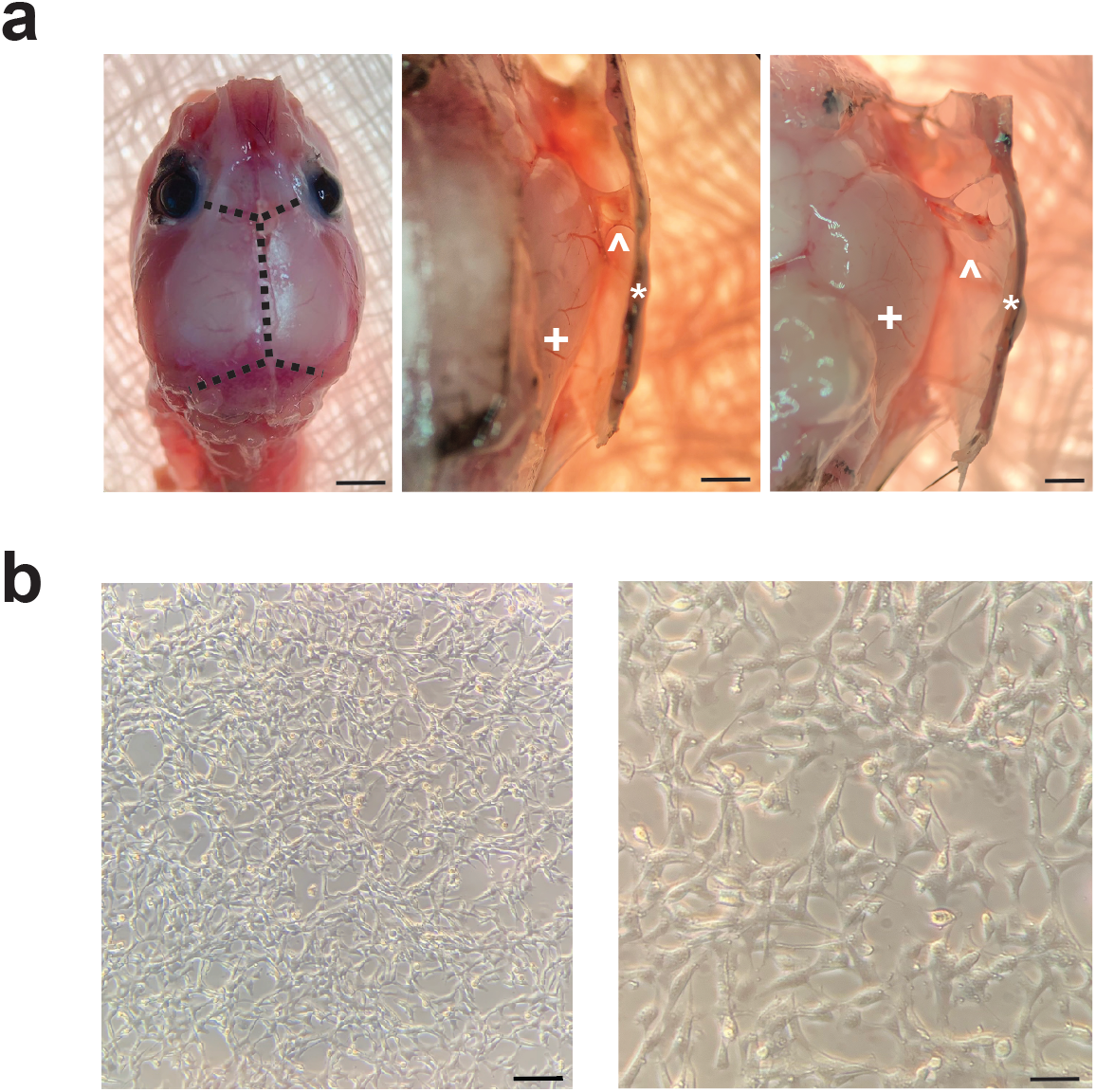
*Pia mater* cells can be isolated and grown in-vitro. a) Incision lines across lambdoid, sagittal and coronal sutures of a mouse skull. *Pia mater* layer (<) as seen under a stereo microscope while skull (*) is being laterally retracted from the brain parenchyma (+). Scale bar, left 5 mm, middle 1 mm, right 1 mm. b) Pial fibroblasts after dissociation using collagenase (200 U/mL) and dispase (1.5 U/mL) and culture in MENCM media at day 7 post-extraction. Scale bar, left 60 um, right 15 um.

**Figure 2:**
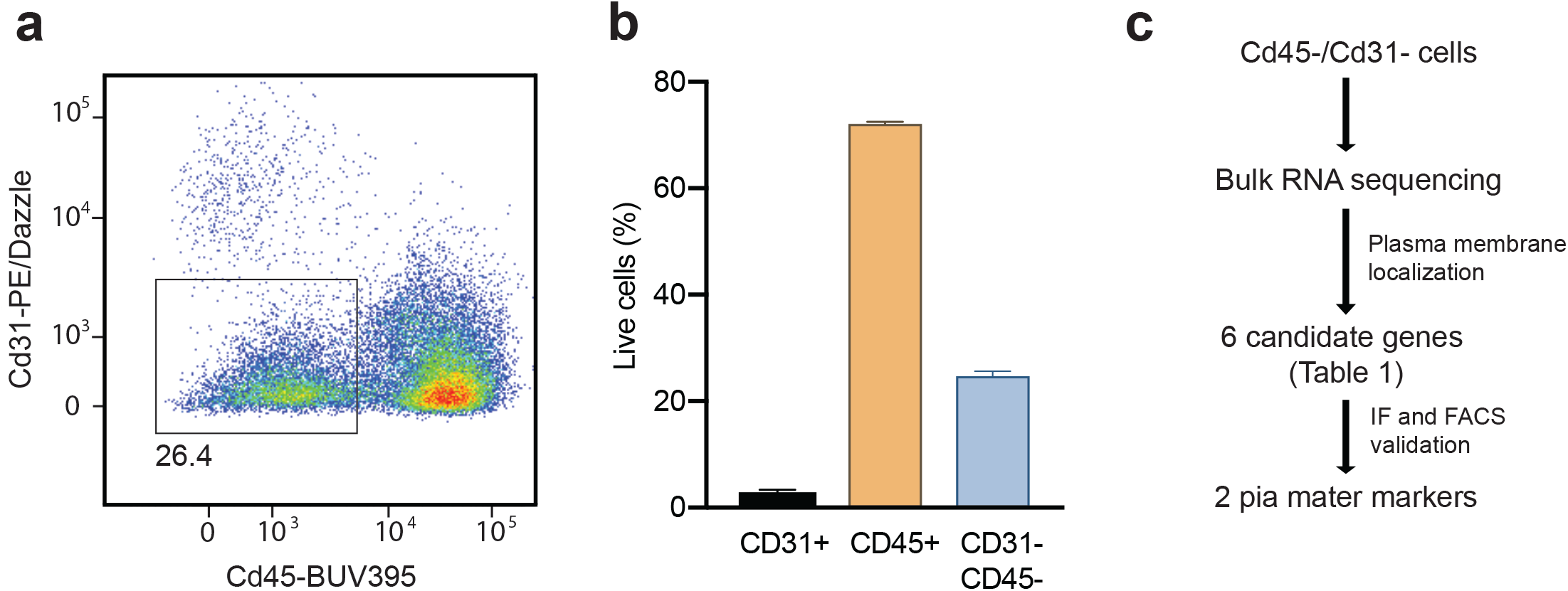
*Pia mater* can be isolated and express unique markers. a) C57Bl/6 mice skulls were dissected as described and *pia mater* tissue was extracted and digested. A single cell suspension was prepared. Erythroid cells were lysed and cells were stained for CD31, CD45 and live/dead green and analyzed by flow cytometry. Cells of interest (*pia mater* cells) were identified by being CD31- and CD45-. b) *Pia mater* cells constituted about 25% of live CD31-/CD45-cells extracted from C57Bl/6 meninges. N=3; All measurements were taken from distinct samples. Error bars represent SE. c) Flow chart describing the methods used for filtering through the bulk RNA sequencing data to identify *pia mater* cell markers.

### Identification of pial cell markers

To identify a cell-surface marker of adult pial cells, CD45^−^CD31^−^ cells from *pia mater* were sorted and subjected to RNA-sequencing. After identifying genes with the highest read counts and filtering for plasma-membrane bound proteins, 6 candidate genes **(Table 1)** were screened with IHC, IF, flow cytometry, or qPCR, complemented with database search (Panther, UniProt). Icam1 and Slc38a2 were further selected as putative pia markers and were validated with a combination of these methods. Flow cytometric analysis showed that 45% of CD457^−^CD3T^−^ cells were Icam1^+^, 30% Slc38a2^+^ and 16% stained for both Icam1 and Slc38a2 (**Figure 3a, Supplementary Figure 3**). Expression of these markers was further quantified in cultured pial cells, showing an increase in both *Icam1* and *Slc38a2* mRNA expression and decrease in *Cd31* expression with passaging. To visualize the identified markers on the pial cell layer in intact adult mouse brains and adult human meninges, laminin was used as a basement membrane marker to locate the overlying pial cell layer. Icam1 and Slc38a2 marked the pial cell monolayer in mouse meninges (**Figure 4a**) and human meninges (**Figure 4b**) overlying the basement membrane. Similarly, a whole-mount staining of pial tissue showed Icam+ and Slc38a2+ cells within the pial layer with cells that co-stained for both markers (**Figure 4c**). As an orthogonal approach, we took advantage of the Allen Brain Map single-cell transcriptomes. We identified five distinct transcriptomic states in a small cluster of mouse vascular-leptomeningeal cells (VLMC), fraction of which is reminiscent of pial fibroblasts. Taking into account mRNA drop-out during single-cell RNA-seq, both *Icam1* and *Slc38a2* mRNA were detected in all of these subsets (**Figure 6a**). The presence of *Icam1* and *Slc38a2* mRNA in the pial tissues was further verified in publicly available RNA in-situ hybridization (ISH) data (**Figure 6b**). To clarify the function of these VLMC cells, we assessed the cellular functions overrepresented within the highly variable genes in both mouse and human VLMC single-cell transcriptomes (**Figure 6c, Supplementary Tables 1, 2**). VLMC cells are in both organisms involved in a surprisingly wide spectrum of physiological functions, including the transport of small molecules, synthesis of extracellular matrix components, and matrix-mediated interactions.

**Figure 3:**
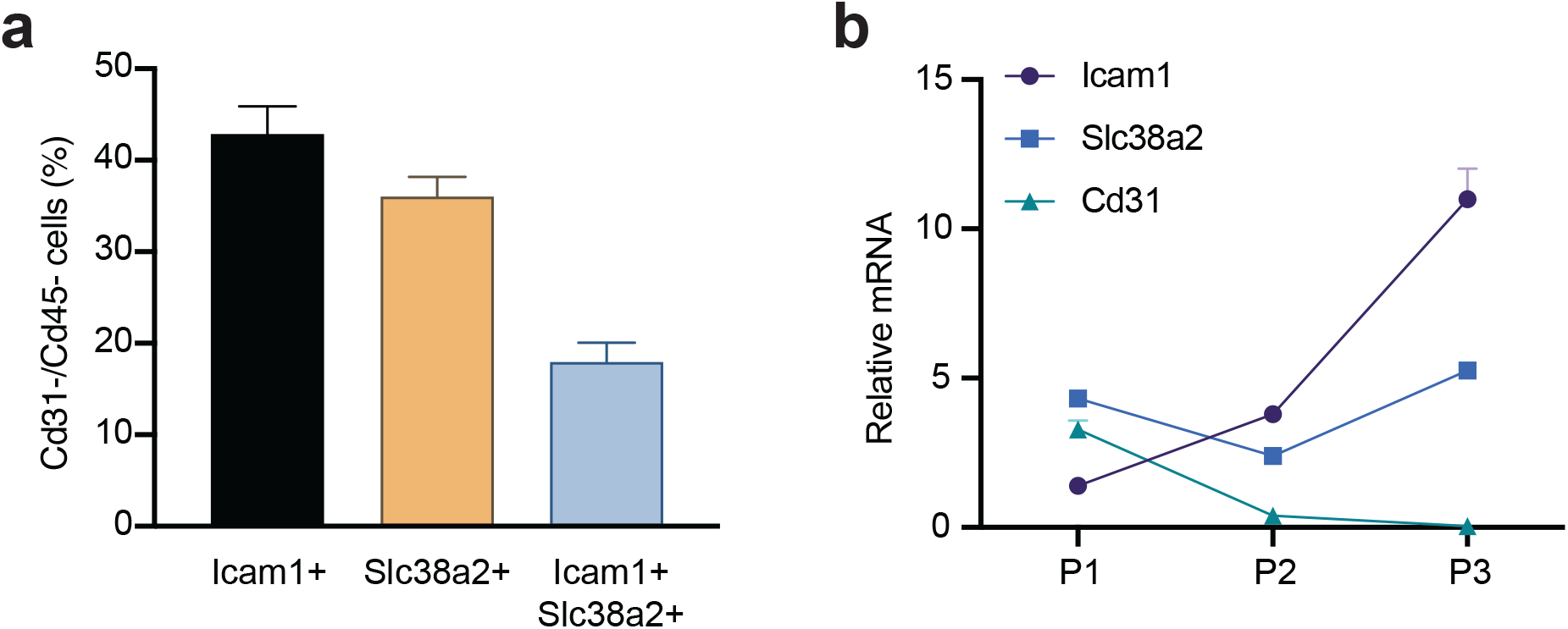
*Pia mater* cells express Icam1 and Slc38a2 and maintain them in culture. a) *Validation of identified pia mater markers on flow cytometry. Pia mater* cells were extracted according to the described protocol. Cells were stained with FITC anti-mouse CD31 (endothelial cells) antibody, FITC anti-mouse CD45 (lymphoid cells) antibody, FITC anti-mouse Ter119 (erythroid cells) antibody and live/dead fixable far red stain. Additionally, cells were either stained for Icam-1, Slc38a2 or both. Error bars represent SE. b) Expression of pia cell markers with passaging of pial cells in vitro. *Pia mater* marker expression was compared to cd31 mRNA expression to ensure selectivity for pial cells in culture. N=3; All measurements were taken from distinct samples. Error bars represent SD.

**Figure 4:**
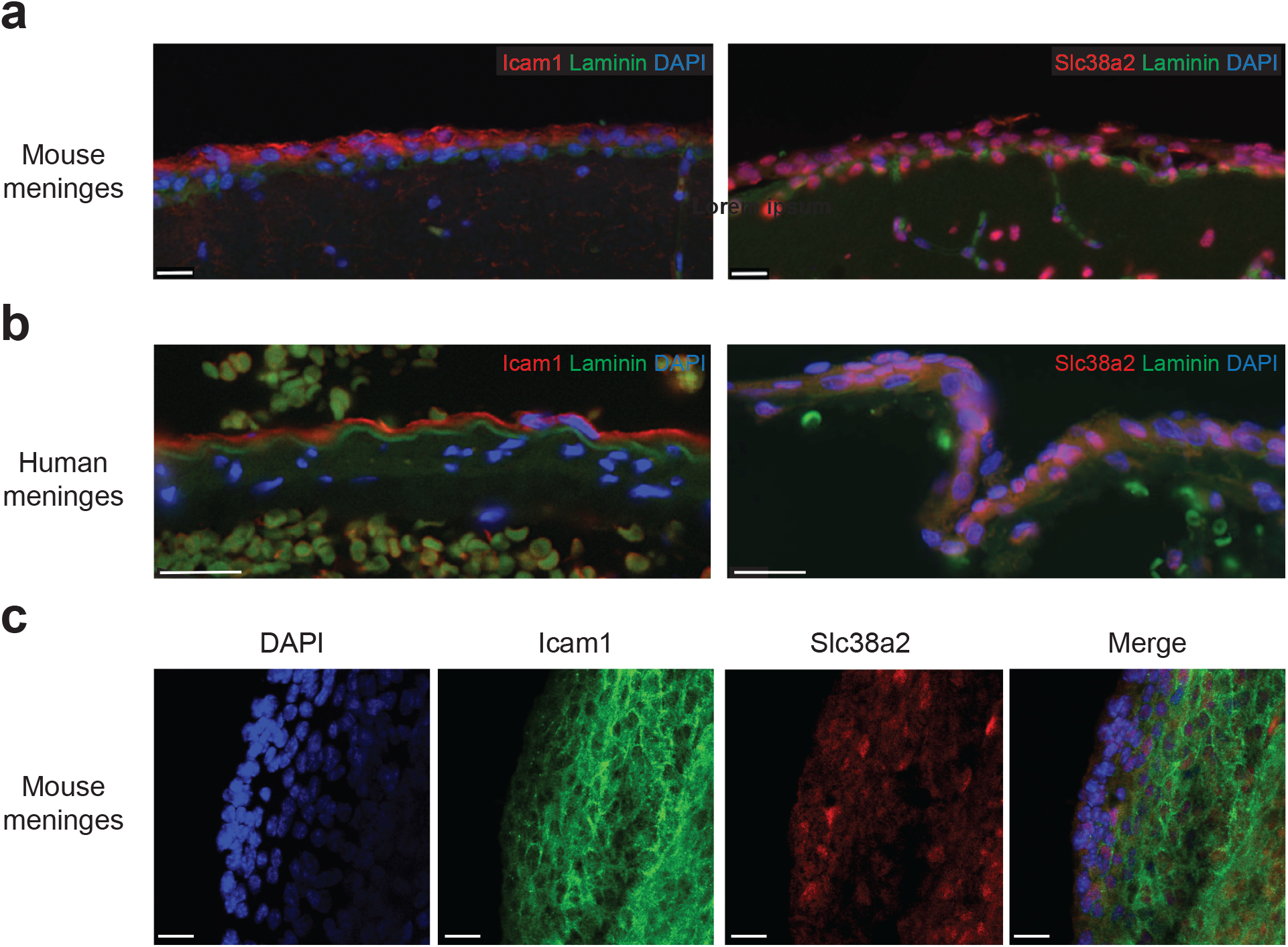
Icam1 and Slc38a2 mark the *pia mater* layer in mice and humans. Visualization of pia mater markers on immunofluorescence in C57Bl/6 mice brains (a), human meninges from autopsy samples (b) and pial tissue whole-mount (c). *Pia mater* cells are visible stained with anti-mouse Slc38a2 antibody or Icam1 antibody (red) overlying laminin (green). DAPI was used for nuclear staining. Scale bar, A, C: 50um, B: 20um. Human FFPE-embedded meninges are unique biological materials that are not available for further distribution.

**Table 1:** Candidate genes identified on RNA sequencing analysis of CD31-/CD45-cells isolated from C57Bl/6 mice pia mater.

| Gene symbol | Gene name |
| --- | --- |
| Pdgfrb | Platelet derived growth factor receptor Beta |
| Ddr2 | Discoidin domain receptor tyrosine kinase 2 |
| Icam1 | Intercellular adhesion molecule 1 |
| Kcne4 | Potassium Voltage-Gated Channel Subfamily E Regulatory Subunit 4 |
| Fcgrt | Fc Fragment Of IgG Receptor And Transporter |
| Slc38a2 | Solute Carrier Family 38 Member 2 |

### Comparison of primary pial fibroblast to fibroblasts isolated from extra-cranial organs

To assess the specificity of these putative pial makers between fibroblasts from other body compartments, we compared them to primary cultures of fibroblasts isolated from lungs and mammary fat pad. Pdgfra was used as a fibroblast marker. While Pdgfra is overexpressed at an mRNA level in lung fibroblasts, protein expression remains equal in all cultured fibroblasts. While Icam1 expression was not exclusive for pial cells, Slc38a2 demonstrated higher expression at both the mRNA and protein levels in pial cells, as compared to lung and MFP fibroblasts (**Figure 5a, b**). Additionally, we also compared the mRNA expression of the identified pial cell markers between pial cells, brain, kidney and spleen tissues using qPCR. Pial tissues reveal a unique Slc38a2 and Icam1 expression signature when compared to other organs (**Supplementary Figure 2**).

**Figure 5:**
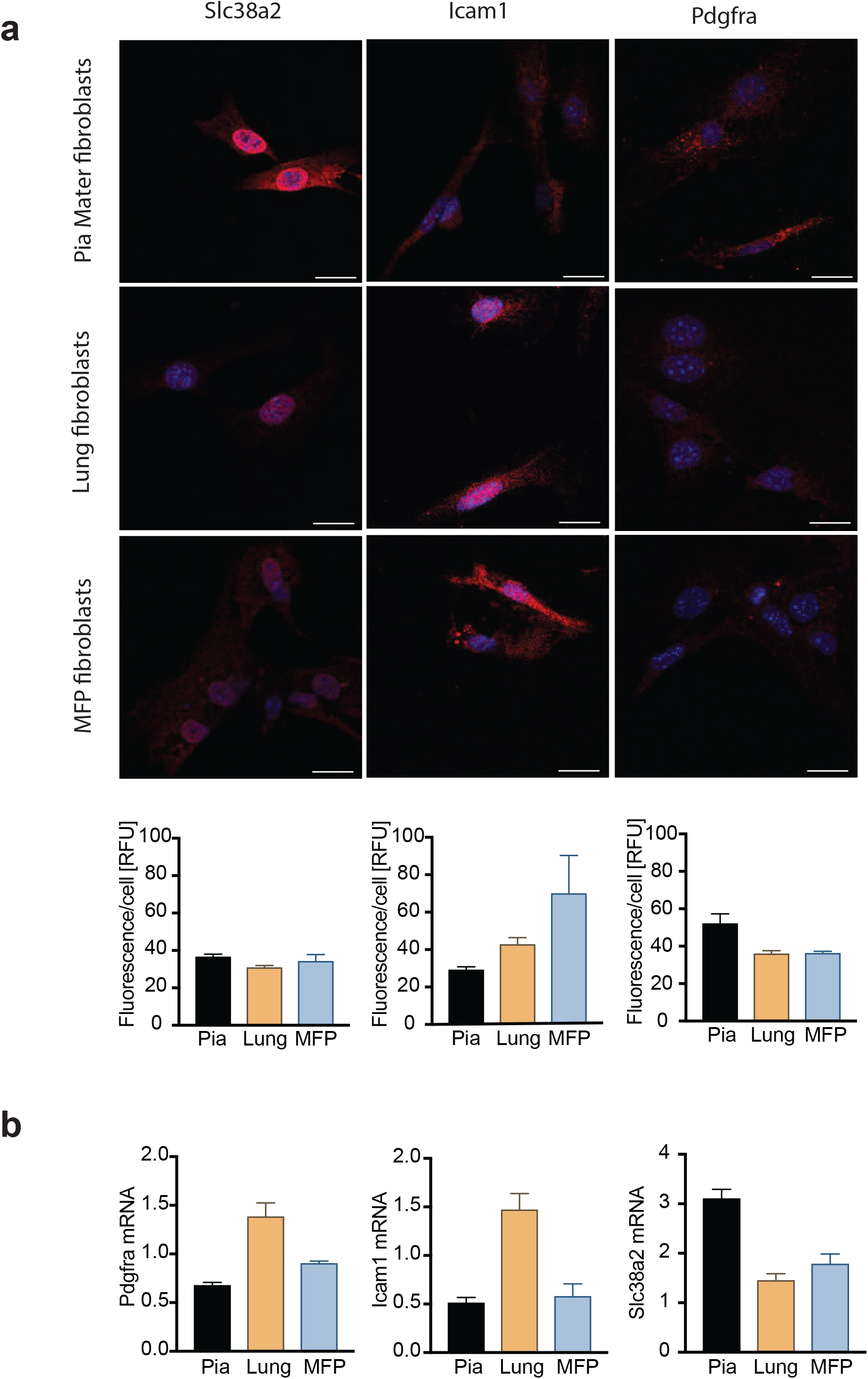
Expression of both Icam1 and Slc38a2 identifies pial cells from other fibroblasts. a) Protein expression of Pdgfra and identified markers (Icam1 and Slc38a2) in *pia mater* cells and fibroblasts from lung and mammary fat pads visualized by immunocytochemistry and quantified using ImageJ (fluorescence intensity/cell area [RFU]). Scale bar, 50um. Data from 3 HPFs per sample; Error bars represent SD. b) mRNA expression of Pdgfra and identified markers (Icam1 and Slc38a2) in *pia mater* cells and fibroblasts from lung and mammary fat pads visualized by qRT-PCR (relative mRNA expression). N=3; All measurements were taken from distinct samples. Error bars represent SD.

**Figure 6:**
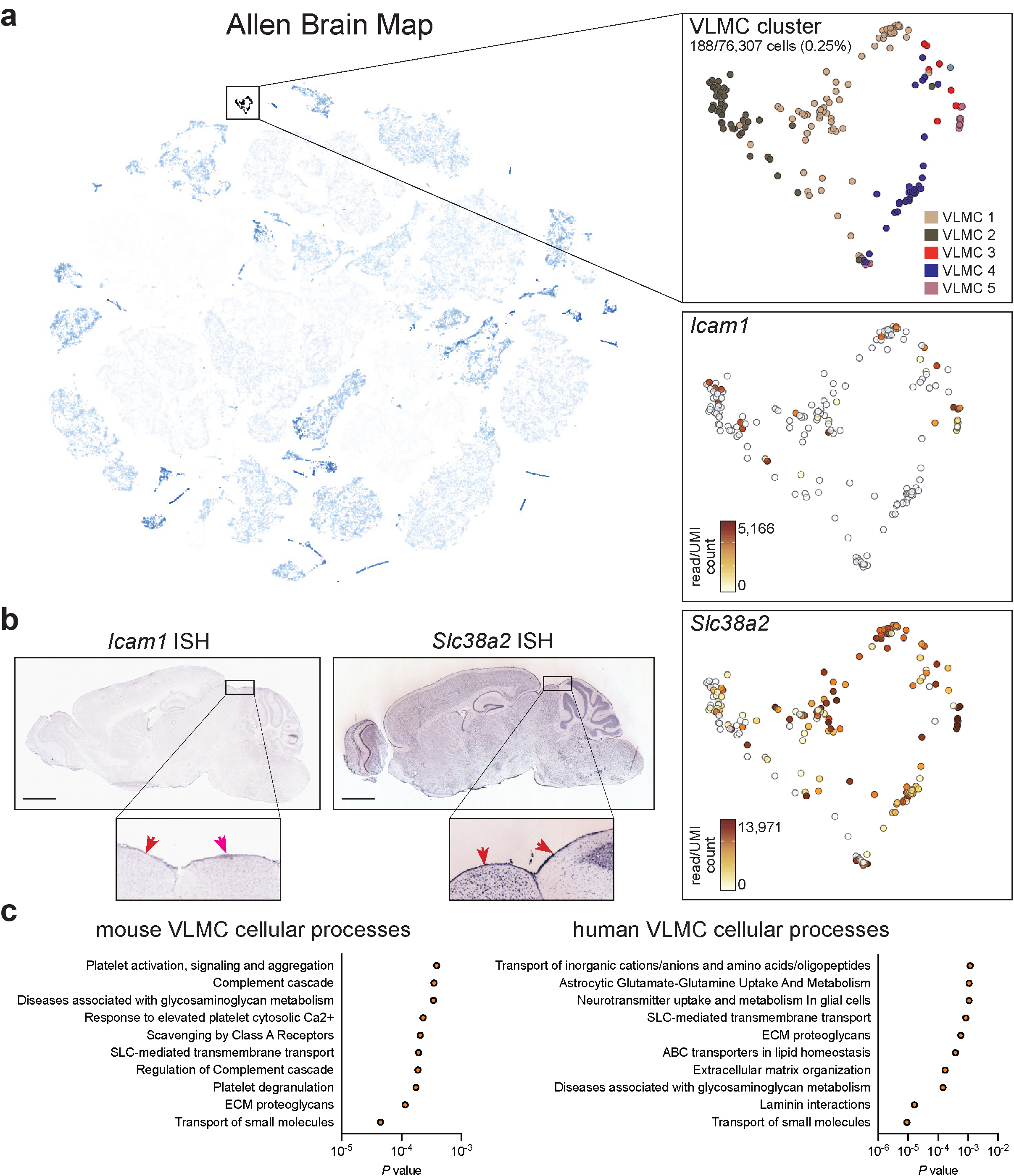
Function of VLMC cells. a) Expression of *Icam1* and *Slc38a2* at single-cell resolution in the cluster of mouse VLMC in Allen Brain Map. b) RNA *in situ* hybridization of *Icam1* and *Slc38a2* in adult mouse brain section. Scale: 1678um. Arrows in inserts point to the *pia mater* layer. c) Functional annotation of highly variable genes in mouse and human VLMC, plot show top ten cellular processes ranked based on *P* value (see also Supplementary Tables 1, 2).

## Discussion

### Applications of the protocol

The structure and function of *pia mater* is not yet well understood and, in general suffers from lack of dedicated study. The presented detailed protocol enables investigators to reliably establish a primary pial cell culture with fibroblast-like characteristics and that retains two putative adult pia markers *in vitro*: Icam1 and Slc38a2. The isolated primary pial cells enable study of conditions that affect leptomeninges, such as infections, autoimmune conditions or cancer. In order to assess the mechanistic and biochemical features of this niche, the primary *pia mater* cells can be isolated, cultured and maintained *in vitro* for a limited number of passages, preserving the characteristic markers described in this manuscript. These markers can be used to purify or identify pial cells for additional downstream applications. Importantly, these markers have a functional role as well, allowing for the interaction of pial cells with other surrounding cells that can be further studied in normal and pathological states.

### Comparison with other methods

Other protocols exist for the bulk isolation of the meninges from mouse or rat brains ^12,13^. However, these protocols fail to distinguish between the different meningeal layers and are usually used to study bulk meningeal immunity or vasculature in response to parenchymal or meningeal diseases. This distinction is of paramount importance. The pachymeninges exist outside the blood brain barriers, are the result of a distinct embryonic lineage, and host a distinct range of pathologies apart from the leptomeninges. Isolation of a clean leptomeningeal population enables dedicated study of this unique anatomic compartment. Using our protocol, researchers can reliably extract and study *pia mater* cells with the help of the identified markers.

### Experimental design

We provide a step-by-step protocol allowing the investigators to specifically dissect mouse *pia mater* tissue that can be further used for a wide variety of downstream applications, including primary cell culture. The major obstacle in this procedure is the identification and gross dissection of *pia mater*. The method in this manuscript provides guidance to the researcher without prior knowledge of mouse meningeal anatomy to reliably isolate this delicate tissue.

Freshly isolated *pia mater* tissue can be digested into single-cell suspension and seeded into cell culture that can be maintained for a limited number of passages and allows to provide mechanistic and biochemical understanding of this tissue. Alternatively, single-cell suspensions can be analyzed with flow cytometry or FACS-purified. Such sorted pial cells from normal mice and mice with meningeal pathologies can be assessed with qPCR or RNA-sequencing to uncover pathology-induced aberrances in signaling pathways or genes-of-interest. Freshly isolated *pia* can be, after brief fixation, used to stain antigens-of-interest in a whole-mount manner.

Using strategy described in this protocol, we identified and validated two putative pial markers, Icam1 and Slc38a2. These markers can be used separately or in tandem to identify the pial layer in *ex vivo* mouse brains or human autopsy specimens.

### Limitations

One of the limitations of this method is the requirement of deep anatomical understanding of murine meninges that is critical for precise *pia mater* identification and extraction. This will ensure maximal efficiency of pial cell recovery without cross-contamination with cells derived from other meningeal layers or brain tissue. To avoid cross-contamination with surrounding tissue, the experimenter will need to carefully peel the pia from the brain parenchyma while laterally retracting the skull. To ensure sufficient viability of pial cells, dissection must occur within 2-3 minutes after euthanasia and tissues must be placed into ice-cold saline. In summary, this technique requires an optimal balance between speed and precision to achieve a successful *pia mater* isolation.

## Materials and Methods

### Biological materials

Mice: All animal studies were approved by the MSKCC Institutional Animal Care and Use Committee under protocol 18-01-002. Female and male C57BL/6J (Jackson Laboratory) mice were housed in the MSKCC vivarium. All mice were at 8-12 weeks of age.

### Reagents

*Media and Reagents*

1. Meningeal Cell Medium (MenCM) (Cat. 1401), supplemented with 1% Penicillin/Streptomycin (Cat. 0503), 2% Fetal Bovine Serum (Cat. 0010), and 1% Meningeal cell growth supplement (Cat. 1452; all ScienCell). CRITICAL: This medium is optimized for leptomeningeal cell cultures and other media do not support culture of primary pial cells.
2. Freezing media: 10% DMSO in FBS
3. Trypsin EDTA (0.25%), Phenol red (Fisher Scientific, Cat. 25-200-056)
4. Purified Collagenase (Worthington Biochemical, Cat. LS005273)
5. Dispase (5 U/mL) (STEMCELL Technologies, Cat. 07913)
6. Phosphate-buffered saline (PBS) 10x
7. Fetal Bovine Serum (FBS) (Thermo Fisher Scientific, Cat. 10437028)

*Immunocytochemistry/Immunofluorescence*

1. 6-well tissue culture plates (Corning, Cat. 353046)
2. Nunc Lab-Tek 4-chamber slide system, sterile (Thermo Scientific, 154526PK)
3. Corning rectangular slide cover glass (Corning, Cat. 2975245)
4. Ice-cold phosphate-buffered saline
5. Dako Target retrieval solution, 10x (Agilent, S1699)
6. Dehydration/rehydration solutions: Xylenes (Sigma Aldrich, Cat. 534056), 100% ethanol, 90% Ethanol, 80% Ethanol, 70% Ethanol (all Fisher Scientific)
7. Wash buffer: Phosphate-buffered saline Tween-20
8. DAPI (Fisher Scientific, Cat. D1306)
9. Confocal microscope (Olympus, FV1000)
10. Goat Icam1 (Cd54) polyclonal anti-mouse antibody (R&D systems Cat. AF796)
11. Goat ICAM1 (CD54) polyclonal anti-human antibody (R&D systems Cat. BBA17)
12. Rabbit Pdgfr-alpha monoclonal antibody (Novus Biologicals Cat. JF104-6)
13. Rabbit Laminin polyclonal Alexa Fluor 488-conjugated antibody (Novus Biologicals Cat. NB300144AF488)
14. Rabbit Slc38a2/Snat2 polyclonal Cy3 conjugated antibody (Bioss Antibodies Cat. Bs12125R-Cy3)
15. Secondary antibodies for immunofluorescence (Jackson Immunoresearch): Donkey anti-goat IgG Alexa Fluor 488-conjugated (705-545-147), donkey anti-rabbit IgG Cy3-conjugated (711-165-152), donkey anti-goat IgG Cy3-conjugated (705-165-147)

*Flow Cytometry*

1. Corning Falcon Test tube with cell strainer snap cap (Fisher Scientific 08-771-23)
2. PE/Dazzle 594 anti-mouse CD31 antibody (BioLegend Cat. 102429)
3. BUV395 Rat anti-mouse CD45 antibody (BD Biosciences Cat. 564279)
4. Live/Dead fixable green dead cell stain kit (Invitrogen Cat. L34969)
5. Live/Dead fixable far red dead cell stain kit (Invitrogen Cat. L34973)
6. FITC anti-mouse CD45 antibody (Biolegend Cat. 103108)
7. FITC anti-mouse CD31 antibody (Biolegend Cat. 102406)
8. FITC anti-mouse Ter119 antibody (Biolegend Cat. 116206)
9. Fixation buffer: 4% PFA in PBS
10. Permeabilization buffer: 0.3% Triton X-100, 1% FBS in PBS
11. Wash buffer: 1% BSA in PBS
12. ACK lysing buffer (Thermo Fisher Scientific A1049201)
13. Rabbit ICAM-1 (CD54) Polyclonal anti-mouse antibody (Invitrogen Cat. PA5-79430)
14. Rabbit SLC38A2 (SNAT2) polyclonal antibody (LSBio Cat. LS-C179270)
15. PE donkey polyclonal anti-rabbit IgG (Biolegend Cat. 406421)

*RNA extraction and qPCR*

1. TaqMan gene expression mouse assays: *Pdgfra* (Mm00440701_m1), *Icam1* (Mm00516023_m1), *Slc38a2* (Mm00628416_m1), *Actb* (Mm00437762_m1)
2. Qiagen RNeasy Plus Micro kit (Qiagen, Cat. 74034)
3. NanoDrop machine
4. High Capacity Reverse transcription kit (Thermo Fisher Scientific, Cat. 4368814)
5. ViiA 7 system (Applied Biosystems)
6. Applied Biosystems TaqMan Advanced Master Mix (Applied Biosystems, Cat. 4444963)

### Equipment

*Dissection Equipment*

1. Dumont #7 Forceps Inox (Roboz Surgical Cat. RS-5047)
2. McPherson-Vannas Micro Dissecting Spring Scissors (Roboz Surgical Cat. RS-5600). CRITICAL: The size of these scissors is ideal for mouse skulls to prevent injury to the underlying parenchyma.
3. Cushing forceps (Roboz Surgical Cat. RS-5284)
4. Sterile polystyrene 100 mm Petri dishes (Fisher Scientific, Cat. FB0875712)
5. Sterile 1.5 mL microcentrifuge tubes (Fisher Scientific, Cat. 21-403-190)
6. Zeiss Stemi 508 Stereo microscope

*General Equipment*

1. Roto-Bot™ Programmable Rotator (Benchmark Scientific)
2. GentleMACS dissociator (Miltenyi Biotec, Cat. 130-093-235)
3. GentleMACS C tubes (Miltenyi Biotec, Cat.130-096-334)
4. 37°C incubator with humidity and gas control to maintain >95% humidity and 5% CO2

*Tissue Culture Equipment*

1. Tissue culture 12-well plates (Fisher Scientific, Cat. 08-772-29)
2. 15- and 50-ml polypropylene centrifuge tubes (Fisher Scientific, Cat. 05-539-12)
3. 10 mL Nunc serological pipettes (Thermo Fisher Scientific, Cat. 170356)

### Step-by-step protocol

#### Isolation of *pia mater* tissue from mice (Timing: ~ 10 min per mouse)

1. Prepare the tissue dissociation solution: 200 U/mL Collagenase and 1.5 U/mL Dispase in 1mL total of serum-free MenCM media. Keep on ice until pial tissue extraction. CRITICAL: Pial tissue should always be kept on ice to maximize live cell yield. The tissue dissociation solution is optimized for fibroblastoid cell dissociation.
2. Euthanize 8-12 weeks old male or female mice according to the approved animal protocol.
3. Sterilize the mouse head with 70% ethanol. Using large scissors, cut through the skin from the back of the head towards the eyes. Detach as much skin and muscles as possible off the mouse’s skull and down the cervical spine approximately to the C7 level.
4. Sever mouse head just above the shoulders (C7).
5. Cut as caudally to the coronal suture to remove the nasal bone while preserving underlying olfactory lobes.
6. Using a posterior approach, slide scissors through the oral cavity and sever muscles connecting the lower jaw. Discard the lower jaw with attaching muscles. CRITICAL: Proceed with this step with caution to prevent puncturing the overlying palate and injuring the brain tissue.
7. Transfer the skull and attached vertebral column into ice-cold PBS for meningeal extraction.
8. Place skull so that ventral aspect of the jaw is facing down. Insert surgical scissors through spinal cord canal at C7, caudally, and cut vertebrae rostrally on dorsal surface adjacent to spinous processes to reach the base of the skull.
9. Cut the skull shallowly along the lambdoid, sagittal and coronal sutures with care taken not to puncture the underlying brain parenchyma **(Figure 1a)**. CRITICAL: Consider using blunt tip scissors to prevent any damage to underlying meninges.
10. Place the skull and attached vertebra in a 100 mm dish with PBS to proceed with dissection under the stereomicroscope.
11. Place the skull on the side that will be dissected to allow visualization of structures beneath the skull when retracted.
12. Using cushing forceps, take hold of the skull from the sagittal suture and open laterally. Pia extraction is accomplished while opening and carefully retracting the skull with slow motion (**Figure 1a**). CRITICAL: Handle the skull with care to prevent disruption of the underlying meninges. Retract the skull very slowly and carefully while extracting tissue from the cortical surface. Make sure not to puncture the brain parenchyma with the scissors to prevent the contamination of the tissue sample with myelinated tissue that complicates downstream applications. Proceed immediately to the next step. *Dissociation of isolated tissue* **(Timing: ~2h 15min)**
13. Extract the transparent tissue adherent to the cerebral cortex using Dumont #7 forceps, starting from the superior sagittal sinus and moving caudally in a medial to lateral fashion. Cerebral meninges must be removed piecewise. CRITICAL: Make sure not to puncture the brain parenchyma with the scissors to prevent the contamination of the tissue sample with myelinated tissue that complicates downstream applications.
14. Place cut tissue immediately in the dissociation solution placed on ice. CAUTION: Pial tissue from 3 mice can be placed in 1mL of the dissociation solution. Do not allow the tissues to dry out.
15. Place tissue on rotator for 2 hours at 37°C (Speed: 25 rpm; Frequency: 1) until complete dissociation. CRITICAL: Time, speed and frequency have been optimized for pial tissue. A change in these parameters can affect the total yield of live pial cells.
16. Using 1 mL pipettes transfer samples to a 5 mL polystyrene tube through a 35 μm cell strainer cap. Do not push too quickly to prevent friction generation by the pipette against the screen.
17. Wash the strainer with additional 3 mL of DME HG to dislodge any cells remaining on/in the strainer. CRITICAL: Keep samples on ice to prevent cell death. Proceed immediately to the next step. Cells can be either cultured (Step 18), analyzed by flow cytometry (Step 18’) or processed for bulk RNA sequencing. *Seeding of isolated tissue* **(Timing: ~ 20 min)**
18. Centrifuge tube at 37°C, 250 g for 5 mins. Discard the supernatant.
19. Resuspend cells in 1 mL of ACK lysing buffer for 5 mins at room temperature to lyse remaining red blood cells.
20. Centrifuge tube at 37°C, 250 g for 5 mins. Discard the supernatant.
21. Resuspend tissue/cells in 1 mL of complete MenCM media and seed in a 35mm tissue culture dish at 37°C.
22. After 48 hrs, carefully aspirate the culture medium and replace with fresh new MenCM medium (**Figure 1b**) and allow cells to grow until 95% confluence is reached. Selectivity for leptomeningeal cells is achieved by using the meningeal specific medium with its meninges-specific supplements. Cells were either assessed for viability with passaging (Step 23) or with repeated freeze-thaw cycles (Step 23’).

#### Identifying extracted sample constituents (Timing: ~ 2 h)

After filtering the extracted sample through the cell strainer (Step 15), cells are stained with PE/Dazzle

594 anti-mouse CD31 (PECAM-1) antibody, BUV395 anti-mouse CD45 antibody and Live/Dead fixable green dye to identify endothelial cells and leukocytes, and to exclude dead cells:

18’. Centrifuge cells for 5 mins, 250 g at 4°C, aspirate the supernatant and add 1 mL PBS.
19’. Add 1 μL of viability dye (Live/Dead green), resuspend the pellet and incubate on ice for 30 mins. CRITICAL: Incubate cells away from any light sources.
20’. Add 1 mL 10% FBS in PBS and spin for 5 mins, 250 g at 4°C.
21’. Discard supernatant; add 5 μL of rat serum in 100 μL of 10% FBS in PBS and incubate on ice for 15 mins.
22’. Add 1 mL 10% FBS in PBS and spin for 5 mins, 250 g at 4°C.
23’. Discard supernatant; add 1 μL of conjugated antibody (CD31, CD45) in 100 μL 10% FBS in PBS for 30 mins. CRITICAL: Incubate cells away from any light sources.
24’. Add 1 mL 10% FBS in PBS and spin for 5 mins, 250 g at 4°C.
25’. Resuspend cells in 400 μL of 10% FBS in PBS and keep samples at 4°C.
26’. Analyze samples using a flow cytometer (**Figure 2a, b**). Cell aggregates and debris were excluded from all analyses based on a dual-parameter dot plot in which the pulse ratio [signal height (y-axis):signal area (x-axis)] was displayed. Dead cells were excluded from analysis by staining with LIVE/DEAD Fixable Dead Cell Stain (various amino reactive dyes; Molecular Probes, Thermo Fisher Scientific).

### Maintenance of *pia mater* cells in culture

Given the labor-intensive methods of fresh *pia mater* isolation, we assessed the maintenance of these cells in culture.

#### 23. Assessment of cell viability with passaging

Pial cells were collected from 3 mice (according to the above protocol) and seeded in a 12-well plate well. 90-95% confluency was reached at day 8 post extraction. Cells were passaged every 4 days after reaching approximately 90-95% confluence into other 12-well plate wells. Total number of cells from all wells were counted after every passage (**Supplementary Figure 1a**).

#### 23’. Assessment of cell viability after freezing

Pial cells were collected from 3 mice (according to the above protocol) and seeded in a 12-well plate. 90-95% confluency was reached at day 8 post extraction. Cells were passaged every 4 days after initial confluence into other 12-well plate wells. Freezing was attempted at passages 2 (F1), 3 (F2) and 4 (F3). Cell viability was assessed for all wells at passage 5 using trypan blue dye **(Supplementary Figure 1b).**

### Identifying markers for *pia mater*

#### Transcriptomic analysis

RNA from FACS-sorted CD3T^−^CD45^−^ cells from freshly isolated, 5 male C57Bl/6 mice-derived pial tissue was extracted using RNeasy Micro Kit (Qiagen, #74034). After RiboGreen quantification and quality control by Agilent BioAnalyzer, 2 ng total RNA with RNA integrity numbers ranging from 9.7 to 10 underwent amplification using the SMART-Seq v4 Ultra Low Input RNA Kit (Clonetech catalog # 63488), with 12 cycles of amplification. Subsequently, 10ng of amplified cDNA was used to prepare libraries with the KAPA Hyper Prep Kit (Kapa Biosystems KK8504) using 8 cycles of PCR. Samples were barcoded and run on a HiSeq 4000 in a 50bp/50bp paired end run, using the HiSeq 3000/4000 SBS Kit (Illumina). An average of 36 million paired reads were generated per sample. Reads from generated FASTQ files were quality checked and mapped to the mouse reference genome (mm10) using STAR2.5.0.a. The expression count matrix of uniquely mapped reads was computed with HTseq v0.5.3. Raw counts were normalized by library size using DESeq pipeline in R v3.6.0 running in RStudio v1.0.143.

To select pial marker(s) to enable further flow cytometric analysis, the first 100 genes with the highest normalized raw counts were analyzed by Panther (www.pantherdb.org) and selected for membrane localization. Genes specific to non-plasma membrane proteins were excluded (**Figure 2c**).

#### Real-time quantitative PCR analysis of gene expression (Timing: ~ 4.5 h)

Total RNA was isolated from samples with the Qiagen RNeasy Plus Micro kit according to standard protocol. RNA concentrations were quantified using a NanoDrop machine. 2 μg of RNA was used for cDNA synthesis using the High Capacity Reverse transcription kit (Thermo Fisher Scientific #4368814). Relative qPCR analysis incorporated one housekeeping gene, *Actb* (Mm00437762_m1). Expression levels of genes of interest were measured using the following TaqMan probes (all Applied Biosystems): *Cdkn2a* (Mm00494449_m1), *Icam1* (Mm00516023_m1), *Slc38a2* (Mm00628416_m1), and *Pdgfra* (Mm00440701_m1). qPCR analysis of gene expression was performed on the ViiA 7 system (Applied Biosystems) with the Applied Biosystems TaqMan Advanced Master Mix (Applied Biosystems #4444963). The experiment was repeated in triplicate. Data were processed using the 2^−ΔΔCq^ method.

#### Validation of pial fibroblast markers on flow cytometry

##### Slc38a2

*Pia mater* cells were isolated according to the above protocol, and stained for CD31 (1:100), CD45 (1:100), Ter119 (1:100; all Biolegend), Live/Dead far red (1:1000; Invitrogen) and Slc38a2 (1:500; LSBio). In order to detect the Slc38a2 signal, PE anti-rabbit was added to the sample before flow cytometric analysis (1:200). Cells were fixed with 4% PFA in PBS and permeabilized using 0.3% Triton-X100 in PBS for 15 min at room temperature (**Figure 3a**).

##### Icam1

*Pia mater* cells were isolated according to the above protocol, and stained for CD31 (1:100), CD45 (1:100), Ter119 (1:100; all Biolegend), Live/Dead far red (1:1000; Invitrogen) and Icam1 (10ug/mL; Invitrogen). In order to detect the Icam1 signal, PE anti-rabbit was added to the sample before flow cytometric analysis (1:200) (**Figure 3a**).

#### Visualization of identified markers on fibroblasts using immunocytochemistry

*Pia mater*, mammary fat pad (MFP) and lung tissues were collected from female C57Bl/6 mice. Mammary fat pads and lung lobes were minced, mechanically dissociated using GentleMACS (Miltenyi Biotec) and digested in a mixture of collagenase (200 U/mL) and dispase (1.5 U/mL) in serum-free MenCM media for 2 h at 37°C. Additionally, MFPs and lung tissue were mechanically dissociated every 20 minutes. Suspension was then washed, filtered through a 70 micron mesh, and seeded complete MenCM media. Pial tissue was dissociated and cultured as previously described. After 24 hrs of culture, cells are washed with PBS and fresh MenCM media is added to pial fibroblasts, lung fibroblasts and MFP fibroblasts. When cells reached the 80% confluency, 1 × 10^3^ cells (pial fibroblasts, lung fibroblasts, MFP fibroblasts) were seeded in 4-Chamber slides (Lab-Tek) and left to attach overnight, then fixed with 4% paraformaldehyde and permeabilized using 0.3% Triton X-100 (Sigma-Aldrich, T8787). The slides were blocked at room temperature for 1 hr in 2.5% normal horse serum in PBS (Vector Laboratories). Samples were incubated for 1hr with primary antibodies: goat anti-Icam1 (1:25), rabbit anti-SLC38A2 (1:100). After 3 washes with TBST, samples were incubated for 1 hr with the appropriate secondary antibodies. Cell nuclei were stained using DAPI (Life Technologies). Images were acquired with a confocal microscope (Olympus, FV1000), and analyzed with ImageJ (NIH).

#### *Validation of pial fibroblast markers with immunofluorescence* (Timing: ~ 6 h)

Tissues were collected as described above. All tissue was fixed in 10% neutral buffered formalin (Sigma–Aldrich, HT501128) overnight at 4°C. The brains were then washed with tap water, sliced into four, 3-4 mm thick coronal sections, placed into a plastic cartridge, and soaked in 70% ethanol. The tissue was then dehydrated though serially increasing concentrations of ethanol and embedded into a paraffin block. 5 μm thick sections were used for staining. Antigen unmasking was done by incubation of sections in Target Retrieval Solution (Dako, S1699) in steamer for 30 min and blocking of unspecific sites was performed with 2.5% horse serum solution (Vector). The primary antibodies were used are as follows: rabbit anti-SLC38A2 (1:100), goat anti-ICAM1 (1:25). For indirect immunofluorescence (IF), samples were incubated with Cy-3 Anti-Goat secondary antibody (1:800) for 30 min. Sections were washed three times in wash buffer (DAKO, S300685), and counter-stained with the nuclear dye DAPI (1:1000) for 5 min, followed by two washes in wash buffer. Sections were transferred onto slides and mounted using Fluoro-Gel mount (Electron Microscopy Sciences, 17985). Images were acquired with a confocal microscope (Olympus, FV1000) and analyzed with ImageJ (NIH).

#### Whole-mount staining of pia mater

Pial tissue was extracted as previously described. Pial tissue was collected in PBS prior to staining. Tissue was fixed with 4% paraformaldehyde and permeabilized using 0.3% Triton X-100 (Sigma-Aldrich, T8787). Pial tissue was blocked at room temperature for 1 hr in 2.5% normal horse serum in PBS (Vector Laboratories). Samples were incubated overnight with primary antibodies at 4C: goat anti-Icam1 (1:25), rabbit anti-SLC38A2 (1:100). After 3 washes with PBST, samples were incubated for 1 hr with AF488 anti-goat secondary antibody (1:800). Cell nuclei were stained using DAPI (Life Technologies). Pial tissue was mounted on a slide prior to imaging. Images were acquired with a confocal microscope (Olympus, FV1000), and analyzed with ImageJ (NIH).

### Troubleshooting

| Step(s) | Problem | Possible reason(s) | Possible solution(s) |
| --- | --- | --- | --- |
| 12, 13 | Difficulty in holding and extracting pial tissue | - Too much liquid in the dish<br>- Forceps with teeth are being used | - Drain as much liquid as possible without completely drying the brain<br>- Use flat forceps to avoid cutting the delicate pial tissue |
| 17 | Low yield of cells after digestion | - Amount of extracted tissue was more than usual<br>- Tissue was not placed on ice for the whole duration of extraction<br>- Tissue was left to dry out<br>- Tissue extraction took more time than usual<br>- Cells lost during centrifugation or aspiration of supernatant | - Add additional dissociation solution if large clumps are still seen after one hour of dissociation<br>- Place tissue on ice for the whole duration of extraction<br>- Time needed for the procedure will decrease with acquired expertise<br>- Aspirate the supernatant from the side opposite to the cell pellet (if visible) |
| 26' | High percentage of dead cells<br>Presence of debris | - Wrong dissociation frequency, speed or time was used.<br>- Dissociation was too rigorous.<br>- Extraction of myelinated tissue with the pial sample | - Use the settings provided in the protocol for optimal dissociation of the pial tissue<br>- Use forceps with narrow tips and extract pial while retracting the skull |

---

**Supplementary Figure 1.**
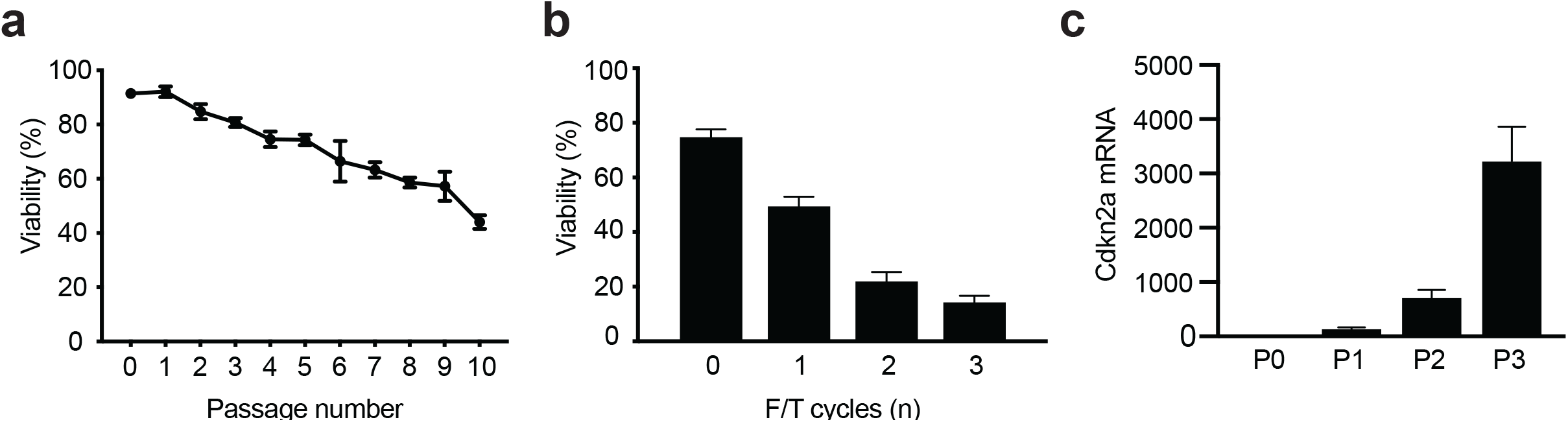
a) *Assessment of cell viability with passaging:* Pial cells were seeded in a 12-well plate well and passaged every 4 days. Total number of cells from all wells were counted after every passage using trypan blue dye. b) *Assessment of cell viability after freezing:* Pial cells were seeded in a 12-well plate and passaged every 4 days; freezing at passages 2 (F1), 3 (F2) and 4 (F3). Cell viability was assessed for all wells at passage 5 using trypan blue dye. c) *Assessment of cellular senescence with passaging:* pial cells were passaged as previously described. mRNA from each the first 3 passages was isolated and Cdkn2a expression levels were determined via qRT-PCR.

**Supplementary Figure 2.**
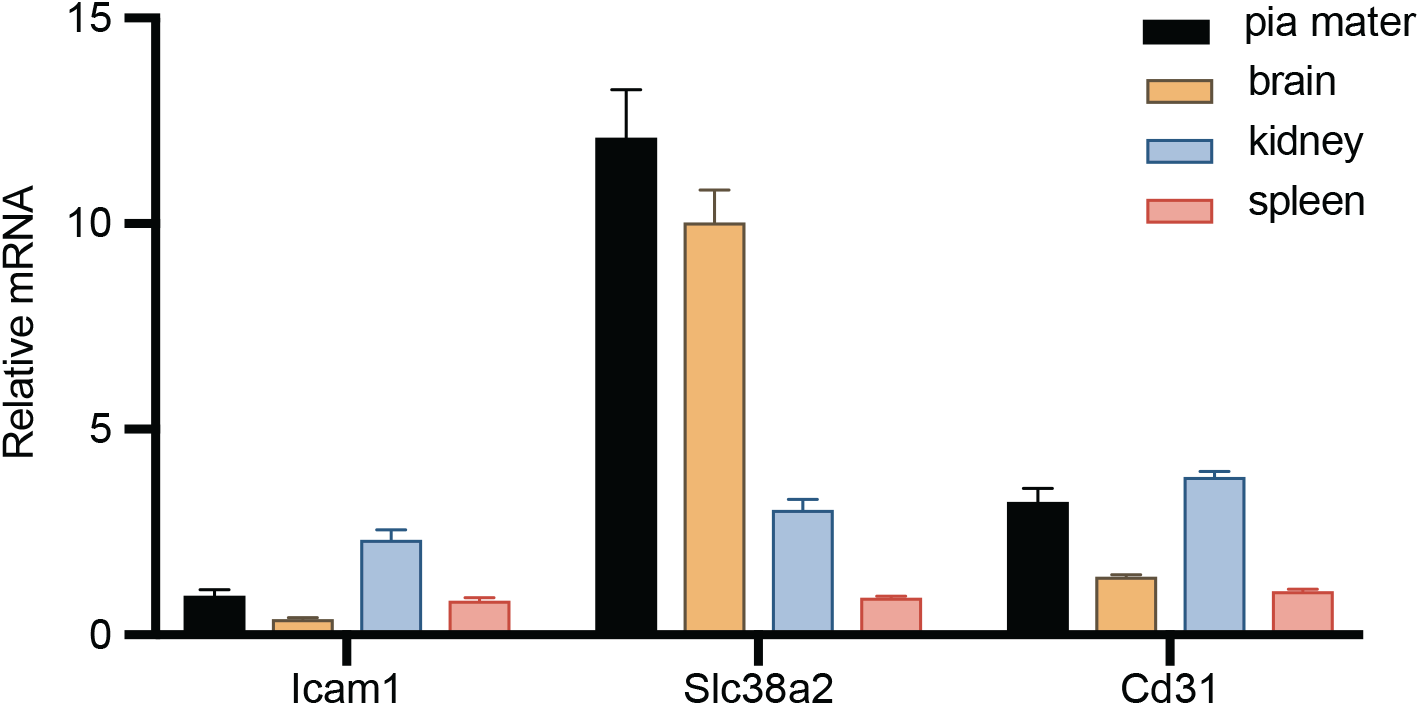
mRNA expression of pia cell markers and cd31 in *pia mater*, brain, spleen and kidney. Total RNA was directly isolated from tissues from these organs and expression of genes of interest was quantified by qRT-PCR (relative mRNA).

**Supplementary Figure 3.**
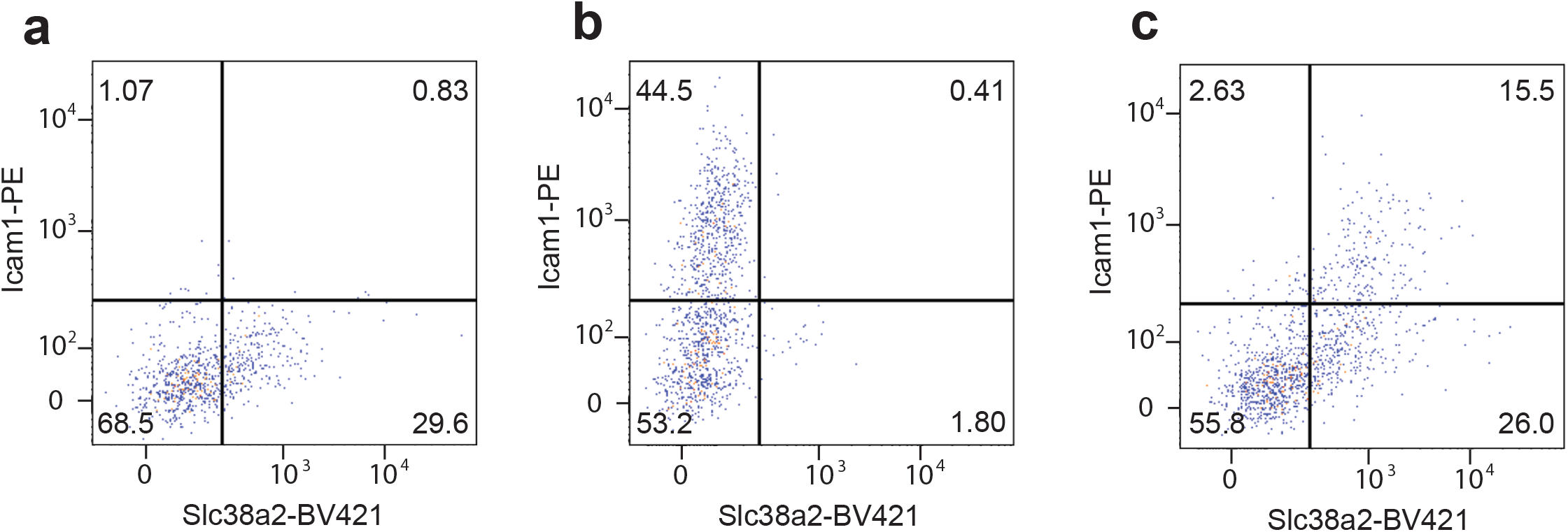
Flow cytometry dot plots for *pia mater* cells co-staining for Icam1 and Slc38a2. a) *Pia mater* cells stained for *Slc38a2*. b) *Pia mater* cells stained for *Icam1*. c) *Pia mater* cells co-stained for *Slc38a2* and *Icam1*.

## Data availability statement

RNA-seq data were deposited to NCBI GEO under the accession number GSE150419. Source data are available from corresponding author upon reasonable request.

## Author contribution statement

A.B. conceived the project and supervised the study. A.B, F.S. and J.R. designed the experiments. F.S., J.R., C.D. and Y.C. performed the experiments. F.S., J.R. and A.B. wrote the manuscript.

## Acknowledgements

We acknowledge the use and help of the Flow Cytometry, Molecular Cytology and Integration Genomics Operation cores at Memorial Sloan Kettering.

---

**Supplementary Table 1:**
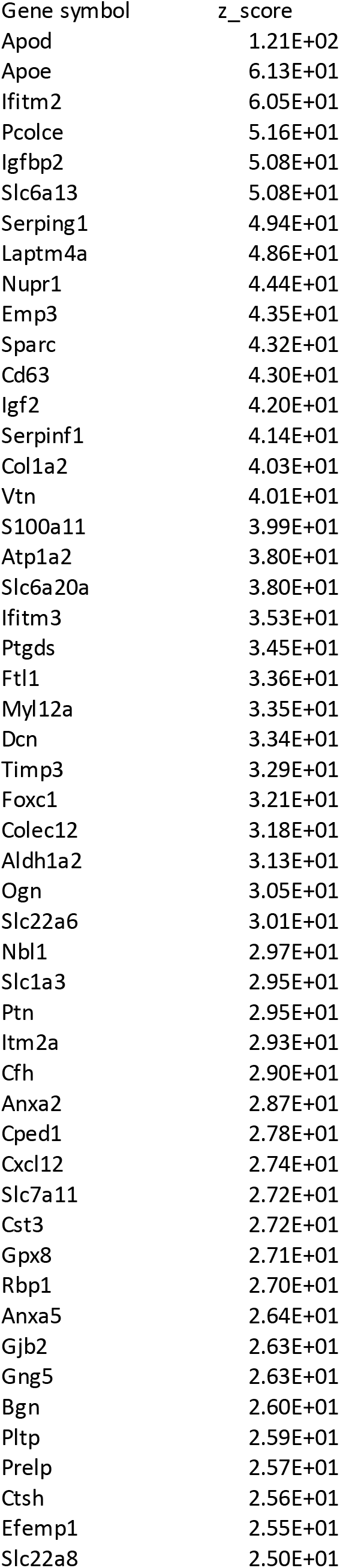
Top 51 highly variable genes in murine VLMC cluster

**Supplementary Table 2:** Top 51 highly variable genes in human VLMC cluster

| Gene symbol | z_score |
| --- | --- |
| ITIH5 | 9.89E+00 |
| ABCA9 | 7.57E+00 |
| DCN | 7.46E+00 |
| ATP1A2 | 6.71E+00 |
| SAT1 | 5.80E+00 |
| NEAT1 | 5.79E+00 |
| SLC7A11 | 5.60E+00 |
| ABCA8 | 5.45E+00 |
| APOD | 5.24E+00 |
| NKD1 | 5.18E+00 |
| PTN | 5.16E+00 |
| FBLN1 | 5.03E+00 |
| LAMC3 | 4.88E+00 |
| PDGFRB | 4.70E+00 |
| CXCL12 | 4.56E+00 |
| AKR1C2 | 4.45E+00 |
| SVIL | 4.34E+00 |
| ABCA6 | 4.19E+00 |
| COLEC12 | 4.14E+00 |
| CYP1B1 | 4.12E+00 |
| UACA | 4.11E+00 |
| AHNAK | 4.04E+00 |
| TPCN1 | 3.95E+00 |
| SLC1A3 | 3.90E+00 |
| ADAM33 | 3.89E+00 |
| LAMA2 | 3.70E+00 |
| COL18A1 | 3.63E+00 |
| CYTL1 | 3.59E+00 |
| SLC6A1 | 3.59E+00 |
| EDN3 | 3.55E+00 |
| ARHGAP29 | 3.47E+00 |
| COL4A5 | 3.47E+00 |
| DAB2 | 3.46E+00 |
| BGN | 3.43E+00 |
| LEPR | 3.41E+00 |
| PTGDS | 3.37E+00 |
| OGN | 3.33E+00 |
| WDR86 | 3.32E+00 |
| PRELP | 3.29E+00 |
| ISLR | 3.26E+00 |
| PDGFRA | 3.26E+00 |
| MRC2 | 3.19E+00 |
| COL15A1 | 3.14E+00 |
| SPRY4 | 3.07E+00 |
| SELENBP1 | 3.04E+00 |
| RHOJ | 3.02E+00 |
| KIAA1755 | 3.01E+00 |
| SASH1 | 2.97E+00 |
| LHFP | 2.92E+00 |
| RPS12P5 | 2.81E+00 |
| KANK2 | 2.70E+00 |

## Notes

### Competing Interest Statement

The authors have declared no competing interest.

